# Kinetochores grip microtubules with directionally asymmetric strength

**DOI:** 10.1101/2024.02.22.581622

**Authors:** Joshua D. Larson, Natalie A. Heitkamp, Lucas E. Murray, Andrew R. Popchock, Sue Biggins, Charles L. Asbury

**Author notes:** Correspondence (J.D.L.), (S.B.), (C.L.A.).

## Abstract

For accurate mitosis, all chromosomes must achieve ‘bi-orientation’, with replicated sister chromatids coupled via kinetochores to the plus ends of opposing microtubules. However, kinetochores first bind the sides of microtubules and subsequently find plus ends by directed transport or when side-attached microtubules shorten and bring their ends to the kinetochores. Mitotic accuracy depends on the selective release of erroneous attachments and proposed mechanisms have focused mainly on plus-end attachments. Whether erroneous side-attachments are distinguished from correct side-attachments is unknown. Here we show that side-attached kinetochores are very sensitive to microtubule polarity, gripping six-fold more strongly when pulled toward plus versus minus ends. This directionally asymmetric grip correlates with changes in the axial arrangement of subcomplexes within the kinetochores, suggesting that internal architecture dictates attachment strength. We propose that the kinetochore’s directional grip promotes accuracy specifically during early mitosis, by stabilizing correct attachments even before both sisters have found plus ends.

## INTRODUCTION

Mitosis is driven by kinetochores, multiprotein complexes that couple chromosomes to the dynamic tips of microtubules. Accurate mitosis requires all chromosomes to become ‘bi-oriented’, with sister chromatids attached via their kinetochores to the plus ends of microtubules emanating from opposite poles of the mitotic spindle. But kinetochores first bind the sides of spindle microtubules^1–4^ and subsequently find plus ends, either when dragged there by their sisters or by plus end-directed motor proteins,^3,5,6^ or when side-attached microtubules shorten, bringing their disassembling plus ends to the kinetochore.^3,7^ Once kinetochores are attached to plus ends *in vivo*, they track persistently with end growth and shortening.^7^ Plus-end tracking powers anaphase, the signature event of mitosis, when sister chromatids are pulled apart.

If sister kinetochores attach correctly to microtubules from opposite spindle poles, they come under tension, and they grip the microtubules stably. Conversely, if they are incorrectly attached, they lack tension and they release to give another chance for proper attachments to form.^8–10^ This selective stabilization of correct, tension-bearing attachments is the fundamental basis for mitotic accuracy. The mechanisms proposed to explain it have focused primarily on plus-end attachments.^9–11^ Aurora B kinase is thought to selectively release plus-end attachments that lack tension,^12^ whereas the higher tension on correct plus-end attachments is thought to protect them from Aurora B.^13^ A catch bond-like behavior can also stabilize tension-bearing plus-end attachments directly, independent of kinase-based regulation.^14,15^ Thus, plus-end attachments are not only key for generating chromosome movements during mitosis, but also for error correction.

Microtubules are protein polymers composed of αβ-tubulin subunits packed together in a head-to-tail fashion, with each subunit oriented identically along the entire length of each filament.^16^ Their structural polarity confers directionality to motor proteins, and it makes the two ends of a microtubule different: β- tubulin is exposed at plus ends, and α-tubulin is exposed at minus ends. Plus ends are faster-growing and more dynamic.^17^ The αβ-tubulin subunits adopt a straight configuration when embedded in the wall of a microtubule,^18^ but at either end of a growing or shortening microtubule, the subunits are often curved.^19,20^ Growing ends are ‘capped’ by newly added subunits that carry GTP, which is hydrolyzed to GDP after assembly into the microtubule wall.^16^ Given how fundamental plus end attachments are for mitosis, kinetochores might possess an intrinsic, preferential affinity specifically for microtubule plus ends. This idea has not been rigorously tested, however, in part because of the difficulty of disentangling the intrinsic properties of kinetochores from other, indirect influences that can affect their behavior *in vivo*.

Kinetochores isolated from budding yeast^14^ are ideal for studying the intrinsic behaviors of kinetochores. Their composition and function are well conserved, they are the simplest known kinetochores (assembling on a single centromeric nucleosome^21,22^ and attaching to a single microtubule^23^), and they are amenable to genetic and biochemical manipulation. The kinetochores of *S. cerevisiae* contain three main types of microtubule-binding elements.^24^ The primary microtubule binder is the Ndc80 complex (Ndc80c), a flexible protein fibril with a globular ‘foot’ (calponin-homology) domain that binds with a stereospecific ‘footprint’ on the outside surface of the microtubule.^25–29^ Multiple Ndc80c fibrils decorate the centromeric nucleosome and recruit numerous Dam1 complexes that strengthen their attachment to the microtubule. (The human Ska complex serves an analogous function.^30^) The Dam1 complexes can assemble into rings encircling the microtubule,^31–33^ potentially organizing the kinetochore into a cage-like arrangement surrounding the plus end.^34,35^ Also recruited to Ndc80c are microtubule-binding TOG-family proteins, such as Stu2^15,36^ (chTOG in humans^37^). Ndc80c itself has no special affinity for the microtubule tip.^38^ However, some evidence suggests that Dam1c might have a preference for the GTP-containing αβ-tubulin subunits that cap growing tips,^31,39^ and Stu2’s TOG domains bind curved αβ-tubulin,^40^ which is found specifically at microtubule ends. Thus, various potential mechanisms could confer a plus-end preference to the kinetochore. Motors can also interact with kinetochores *in vivo* and actively transport them along the sides of microtubules toward the ends.^3,5,6^

Laser trapping has enabled direct studies of the coupling between isolated kinetochores and dynamic microtubule tips *in vitro*.^13–15^ Native kinetochore particles are first purified and then linked to polystyrene microbeads. The kinetochores often bind initially to the sides of individual microtubules, and laser trap tension can then be applied to drag them to the plus ends. Once at plus ends, the kinetochores track persistently with plus end growth and shortening, mimicking their behavior *in vivo*. Notably, however, tracking a growing plus end *in vitro* requires continuous external tension. If tension is removed (e.g., by shuttering the laser trap), a kinetochore-decorated bead usually ‘parks’ on the microtubule wall, staying attached to a fixed point on the lattice while the plus end grows past it. This requirement for external tension suggests that kinetochores might lack any intrinsic plus end preference and that their persistent tracking of plus end growth *in vivo* might depend strictly on the tension exerted by their sisters on the other side of the bipolar spindle, or on interactions with plus end-directed motors. Alternatively, an intrinsic plus end preference might exist but escape detection, either because an effective test has not yet been devised, or because during kinetochore purification a component required for the preference is inadvertently lost.

Here we show that isolated kinetochores do possess a strong, intrinsic preference for plus ends. Single molecule fluorescence microscopy reveals that individual kinetochores assembled *de novo* in whole cell extracts capture microtubules overwhelmingly by their plus-ends. Laser trap experiments show that native kinetochore particles attach dynamic microtubule tips with a substantially higher strength at plus ends than at minus ends. Strikingly, the kinetochores also grip the sides of microtubules with highly direction-dependent strength, indicating an intrinsic sensitivity to the structural polarity of the microtubule wall. Sub-diffraction localization of fluorescent kinetochore subcomplexes indicates that plus end-attached kinetochores are organized with DNA- and microtubule-binding elements separated along the microtubule axis, matching the physiological arrangement during metaphase. However, side-attached kinetochores adopt a more compact axial arrangement specifically when they are pulled toward a minus end. These observations suggest that both the plus-end preference and the directionally asymmetric grip of the kinetochore arise from its asymmetric architecture. The kinetochore’s asymmetric grip is similar to the directional binding of actin filaments by vinculin,^41^ talin,^42^ and α-catenin,^43^ behaviors which are thought to stabilize attachments between focal adhesions and appropriately oriented F-actin.^44,45^ We propose that the directionally asymmetric grip of the kinetochore stabilizes its attachment to correctly oriented microtubules specifically during early mitosis, even before both sisters have found plus ends.

## RESULTS

### Individual kinetochores including outer microtubule-binding subcomplexes assemble *de novo*

We recently showed that assembly of kinetochores *de novo* in yeast cell lysates can be directly observed at the single molecule level using total internal reflection fluorescence (TIRF) microscopy.^46^ Our approach revealed molecular requirements for the wrapping of centromeric DNA around the centromere-specific histone H3 variant, Cse4 (CENP-A), which creates the chromosomal foundation for the kinetochore. However, the extent of recruitment of microtubule-binding kinetochore elements and their functional attachment to microtubules remained unexplored.

To measure the recruitment of microtubule-binding elements onto single centromeric DNAs, we tethered 180-bp Atto565-labeled DNAs sparsely onto passivated coverslip surfaces through avidin-biotin linkages. The surface-tethered centromeric DNAs were then incubated for an hour with lysate from a yeast strain harboring an endogenous GFP tag fused to the C-terminus of Ndc80 (Figure 1A). After incubation, the lysate was washed away, multi-wavelength TIRF images were collected, and colocalization single molecule spectroscopy (CoSMoS) was performed (Figure 1B).^46–49^ The fraction of wild type centromeric DNA molecules that were decorated with Ndc80-GFP was 3.8 ± 2.3% (Figure 1C). Analysis of photobleaching suggested that many of these assemblies carried multiple copies of Ndc80-GFP (Figure S2). We also imaged assemblies after incubation with lysates from strains harboring the inner-kinetochore proteins, Ndc10-GFP (part of the DNA-binding Cbf3 complex), Cse4-GFP (part of the centromeric nucleosome), and Ctf19-GFP (part of the constitutive centromere-associated network, or CCAN) (Figures 1A and 1B). As we previously reported,^46^ *de novo* assembly of the inner kinetochore was highly efficient, with 28 ± 1%, 40 ± 1%, and 12 ± 1% of centromeric DNAs recruiting Ndc10-GFP, Cse4-GFP, and Ctf19-GFP, respectively (Figure 1C). In addition to the wild type DNA construct, we tested assembly on a negative control mutant DNA (Figure S1) containing a three base pair substitution that eliminates centromere function *in vivo*^50–52^ and *in vitro*.^53^ No more than 0.14% of these negative control mutant DNAs colocalized with GFP-tagged kinetochore components (Figure S1). These observations confirm the specificity of our single molecule kinetochore assembly assay and demonstrate that about one in twenty-five centromeric DNAs recruit microtubule-binding elements. Considering that many hundreds of individual DNAs can be observed in a single field of view, this level of efficiency was sufficient to enable functional, microtubule-binding behaviors of the individual kinetochore assemblies to be studied.

**Figure 1.**
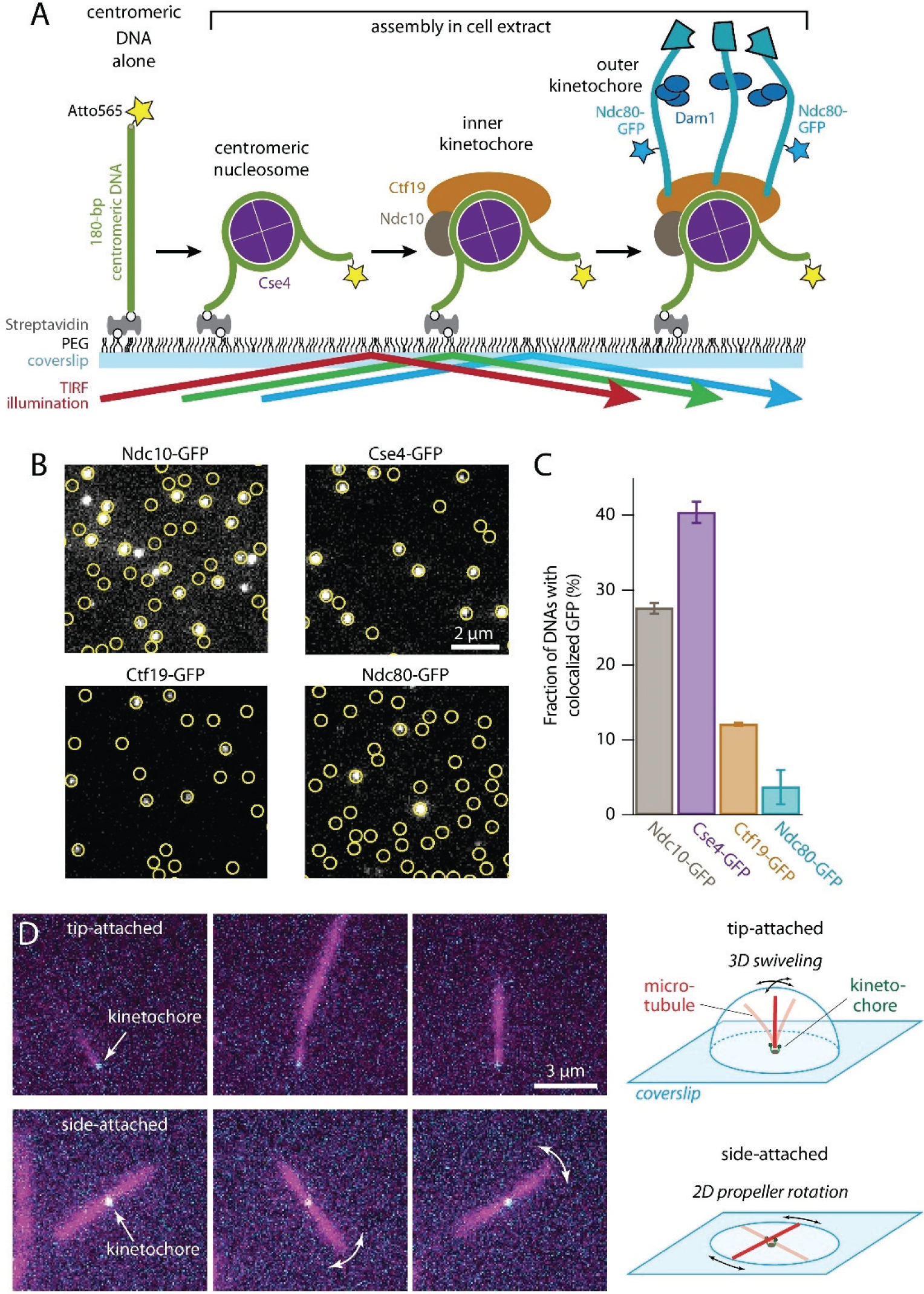
Individual kinetochores assembled *de novo* onto centromeric DNAs capture microtubules. **A)** Schematic of the *in vitro* kinetochore assembly assay. Individual Atto565-labeled centromeric DNAs were tethered sparsely onto a polyethylene glycol (PEG) passivated coverslip surface through biotin-avidin linkages. The surface-tethered DNAs were then incubated for 60 min with yeast whole cell lysate derived from strains with GFP-tagged kinetochore components (Ndc10, Cse4, Ctf19, or Ndc80). Kinetochores assembled spontaneously onto the centromeric DNAs and were then imaged with total internal reflection fluorescence (TIRF) microscopy after washing out the extract. **B)** Kinetochore subcomplexes colocalized with wild type centromeric DNAs. Locations of Atto565-labeled centromeric DNAs (yellow circles) were mapped onto images of GFP-tagged kinetochore subcomplexes (white spots). Scale bar, 2 µm. **C)** Percentages of centromeric DNAs that colocalized with a GFP signal from indicated kinetochore proteins. Bars show average colocalization ± SEM calculated from N > 3,400 DNAs for each kinetochore component from at least 9 fields of view recorded across three independent experiments. **D)** Assembled Ndc80-GFP kinetochores (cyan) readily captured Alexa647 microtubules (magenta) by their tips (top row of images), and sometimes by their sides (bottom row). Tip-captured and side-captured microtubules were easily distinguished by the relative locations of the kinetochore GFP spots and by the Brownian movement of the filaments. The distal ends of tip-captured microtubules swiveled freely in three dimensions (3D), exploring a hemispherical space above the coverslip. Side-captured microtubules mainly rotated in a two-dimensional (2D) plane parallel to the coverslip, in a propeller-like fashion.

### Assembled kinetochores capture microtubules with a strong preference for plus ends

To test for microtubule-binding activity we introduced taxol-stabilized Alexa Fluor 647-labeled microtubules after kinetochore assembly in lysate (Figure 1D). The individual assembled kinetochores readily captured single microtubules. Capture was specific to the kinetochores and did not occur in negative controls with non-functional mutant centromeric DNAs. The kinetochores often captured microtubules by their tips (Figure 1D, Video S1), which colocalized with the fluorescence from Ndc80-GFP. In this tip-attached arrangement, the distal ends of the microtubules swiveled freely by Brownian motion, exploring a hemispherical space above the coverslip. Some kinetochores captured microtubules by their sides. The Brownian movement of these side-captured microtubules was more restricted. Rotation occurred mainly in a plane parallel to the coverslip, swiveling in a propeller-like fashion (Figure 1D, Video S2) with Ndc80-GFP located at the axis of rotation. These observations show that individual *de novo* assembled kinetochores can capture the tips and the sides of microtubules, and they demonstrate flexibility in the tethering of the kinetochores to the coverslip.

To test for preferential plus-end binding, we generated polarity-marked GMPcPP-stabilized microtubules by growing dimly fluorescent plus-end extensions from brightly fluorescent seeds.^54,55^ We then assembled surface-tethered kinetochores, incubated them with the polarity-marked microtubules, and quantified the fraction of microtubules that were captured by plus versus minus ends (Figure 2A). For clear viewing, we applied a gentle flow of buffer (0.6 mL·min^-1^) to keep the kinetochore-attached microtubules parallel to the coverslip and in the plane of focus. More than 82% of tip-bound microtubules (162 out of 196 microtubules examined across eight technical replicates) were captured by their plus ends (Figures 2A and S3). This preference was observed in the absence of motor proteins,^53^ ATP, and microtubule dynamics, implying that kinetochores themselves have a strong intrinsic affinity for features specific to microtubule plus ends.

**Figure 2.**
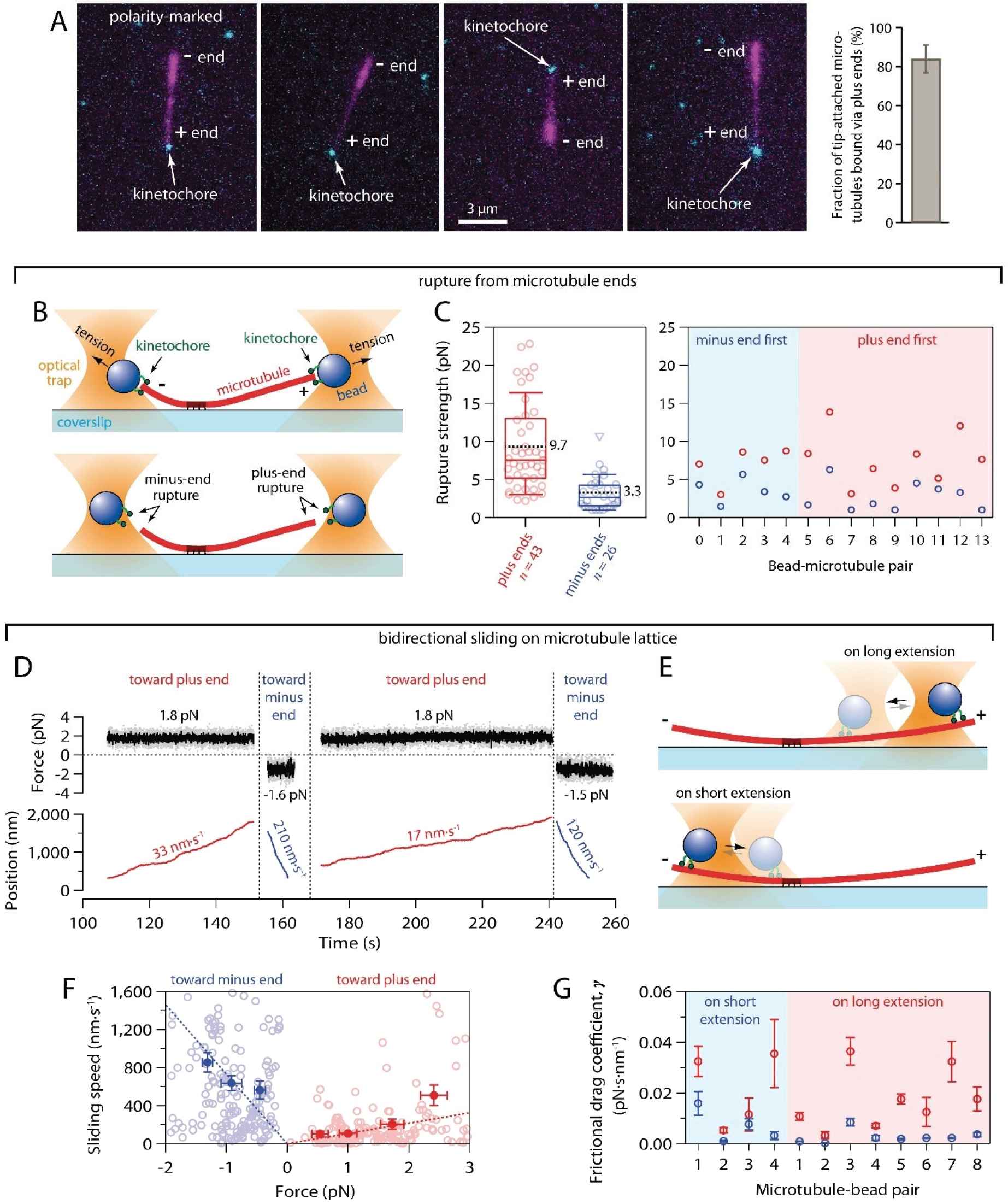
Kinetochores specifically capture plus ends and grip more strongly when pulled toward plus ends. **A)** Kinetochores assembled *de novo* have a strong intrinsic preference for plus ends. Tip-captured, polarity-marked microtubules (magenta), polymerized with dim plus ends and bright minus ends, nearly always bound assembled Ndc80-GFP kinetochores (cyan) by their plus ends. Bar graph shows percentage of tip-attached microtubules that were bound by their plus ends (mean ± SDEV from *N* = 8 experiments examining a total of 196 tip-captured microtubules). **B)** Schematic of the rupture force assay. Native kinetochore particles isolated from yeast were conjugated sparsely to polystyrene microbeads. A laser trap was used to attach a kinetochore-bead to either the plus or minus end of an individual dynamic microtubule and then to measure the rupture strength of the attachment. **C)** *Left:* Distribution of rupture strengths for attachments to plus and minus ends. Open circles represent individual rupture strength measurements. Triangles show censored data, when rupture strength exceeded the maximum force of the trap, or the microtubule broke away from the coverslip surface. Boxes extend from the first to the third quartiles with medians indicated by the central horizontal solid lines. Whiskers extend ± one SDEV from the means, which are indicated by dashed black lines. *Right:* Rupture strengths for individual kinetochore-beads measured sequentially at both ends of the same microtubule, either minus end first or plus end first as indicated. **D)** Measurement of bidirectional sliding friction. Record of force and position versus time for a kinetochore-bead attached to the side of a coverslip-anchored microtubule and pulled alternately toward the plus (red trace) and minus end (blue trace). Additional records are shown in Supplementary Figure 2s2. **E)** Schematic of the sliding assay. Kinetochore-beads were tested on both long and short extensions, to confirm that the speed differential arises from microtubule polarity rather than asymmetric anchorage of the microtubule to the coverslip. **F)** Bidirectional sliding speeds plotted against applied force. Open circles show speeds from individual events. Dotted lines represent fits to these individual speeds. Closed circles represent mean speeds ± SEM after binning by force (*N* = 23 – 81 events per bin). **G)** Frictional drag coefficients for individual kinetochore-beads sliding toward plus (red symbols) and minus ends (blue symbols) on short and long microtubule extensions, as indicated. Beads 1 through 4 were measured sequentially on both extensions of the same microtubule. Symbols represent mean frictional drag coefficient ± SEM (from *N* > 5 events per bead-microtubule pair).

### Plus end attachments support more tension than minus end attachments

Kinetochores sustain tension almost continuously once they are properly end-attached *in vivo*,^56^ so their load-bearing capacity is important for function. We therefore wondered whether the plus-end binding preference uncovered in our TIRF experiments would affect a kinetochore’s load-bearing capacity. Using a laser trap, we measured the rupture strengths of native kinetochore particles isolated from yeast lysate via FLAG immunoprecipitation.^14^ As in our prior work, the native kinetochores were conjugated sparsely to polystyrene microbeads and then attached near the tips of individual dynamic microtubules growing from coverslip-anchored seeds. After an initial preload tension of ∼1 pN was used to slide a kinetochore-bead to the end of a microtubule, the pulling force was gradually increased (at 0.25 pN·s^-1^) until the kinetochore detached from the microtubule (Figure 2B). Plus and minus ends were readily identifiable in these experiments because the plus ends grew faster, extending farther from the coverslip anchored seeds than the minus ends. The distribution of rupture strengths measured at plus ends was very similar to our previous measurements,^14,15^ with a mean strength of 9.7 ± 1.0 pN (mean ± SEM from *N* = 43 plus-end attachments) (Figure 2C, left). Strengths measured at minus ends were substantially weaker, with a three-fold lower mean strength of only 3.3 ± 0.5 pN (*N* = 26 minus-end attachments; *p* = 2×10^-5^, based on a K-S test). In many instances, it was possible to sequentially measure the rupture strength of a single kinetochore-decorated bead at both ends of a microtubule. Irrespective of the order of these measurements, minus-end first or plus-end first, the strength was always higher at the plus end (Figure 2C, right). This control confirms the differential strength and also the reliability of our identification of plus versus minus ends. Altogether, the rupture strength measurements show that attachments of kinetochores to microtubule plus ends can bear substantially more tension than attachments to minus ends.

### Kinetochores grip microtubule sides with direction-dependent strength

Before achieving proper plus end attachments *in vivo*, kinetochores initially bind the sides of microtubules.^1–4^ This physiological behavior is also seen in our laser trap assays with isolated kinetochores. Modest amounts of laser trap tension, 0.5 to 3 pN, will cause these side-attached kinetochores to slide toward the ends. Often, it is easier to detach side-attached kinetochores from microtubules by sliding them toward minus ends. This observation, together with the large strength difference between plus versus minus end attachments, led us to hypothesize that side-attachments might exhibit a polarity preference. To test this idea we used laser trapping to measure the friction between side-attached kinetochores and microtubules.

Microbeads decorated sparsely with native kinetochores were attached to the sides of individual, dynamic microtubules, growing (as described above) from coverslip-anchored seeds. Constant pulling forces between 0.5 and 3 pN were applied parallel to the microtubule axis, and the speeds at which the kinetochore-decorated beads slid along the microtubule were quantified. The direction of force was periodically reversed, to assess friction in both directions relative to microtubule polarity (Figures 2D, 2E, and S4). For every bead-microtubule pair examined, sliding toward the plus end was markedly slower than toward the minus end. Under 1 pN of laser trap tension, the average sliding speeds toward plus versus minus ends differed six-fold (109 ± 11 nm·s^-1^ versus 635 ± 79 nm·s^-1^; mean ± SEM from *N* = 75 and 81 sliding events, respectively, across 24 microtubule-bead pairs) (Figure 2F). Sliding speeds generally increased with force, as expected for protein friction.^57^ To compare friction across many bead-microtubule pairs, measured at different forces and on plus- and minus-end extensions, we divided the applied force, *F*, by the mean sliding speed, *v*, to compute a frictional drag coefficient, *γ* = *F* · *v*^-1^. The frictional drag during plus end-directed sliding was uniformly higher, independent of whether beads were tested on the shorter microtubule extensions (where plus end-directed sliding was toward the coverslip-anchored seeds), or on the longer extensions (where plus end-directed sliding was toward free microtubule ends) (Figure 2G). This control confirms that the speed difference arises from microtubule polarity, rather than from asymmetric anchorage of the microtubule to the coverslip. Altogether, these observations indicate that kinetochores grip the sides of microtubules with a strength that differs markedly depending on the direction of applied force relative to the polarity of the microtubule substrate.

### Assembled kinetochores recapitulate *in vivo* architecture when attached to microtubule plus ends

To understand the basis for direction-sensitivity, we sought to examine the architecture of the isolated kinetochores. When kinetochores are properly attached to microtubule plus ends *in vivo*, their molecular components are spatially organized. Fibrillar Ndc80 complexes align with the microtubule axis, DNA-binding ‘inner’ kinetochore elements are oriented toward plus ends, and ‘outer’ microtubule-binding elements project toward minus ends and spindle poles.^58–60^ To examine the configuration of kinetochores assembled *de novo*, we mapped the relative positions of specific fluorescent-tagged kinetochore components along the microtubule axis by locating their centers of fluorescence. Ndc80-GFP kinetochores were assembled onto Atto565-labeled DNAs and exposed to taxol-stabilized, Alexa Fluor 647-labeled microtubules. A gentle flow of buffer (0.6 mL·min^-1^) was applied with a syringe pump to exert sub-piconewton viscous forces that aligned kinetochore-attached microtubules with the plane of the coverslip (Figure 3A). We oscillated the flow direction, causing the microtubules to flip back and forth, reorienting their long axes by 180° with each reversal of the flow (Figures 3B and 3C). The attached kinetochores also reoriented together with the microtubules, allowing us to measure distances from specific fluorescent-tagged kinetochore components to the tether point on the coverslip surface with nanometer accuracy. Initially, we focused on kinetochores that had captured microtubules by their ends.

**Figure 3.**
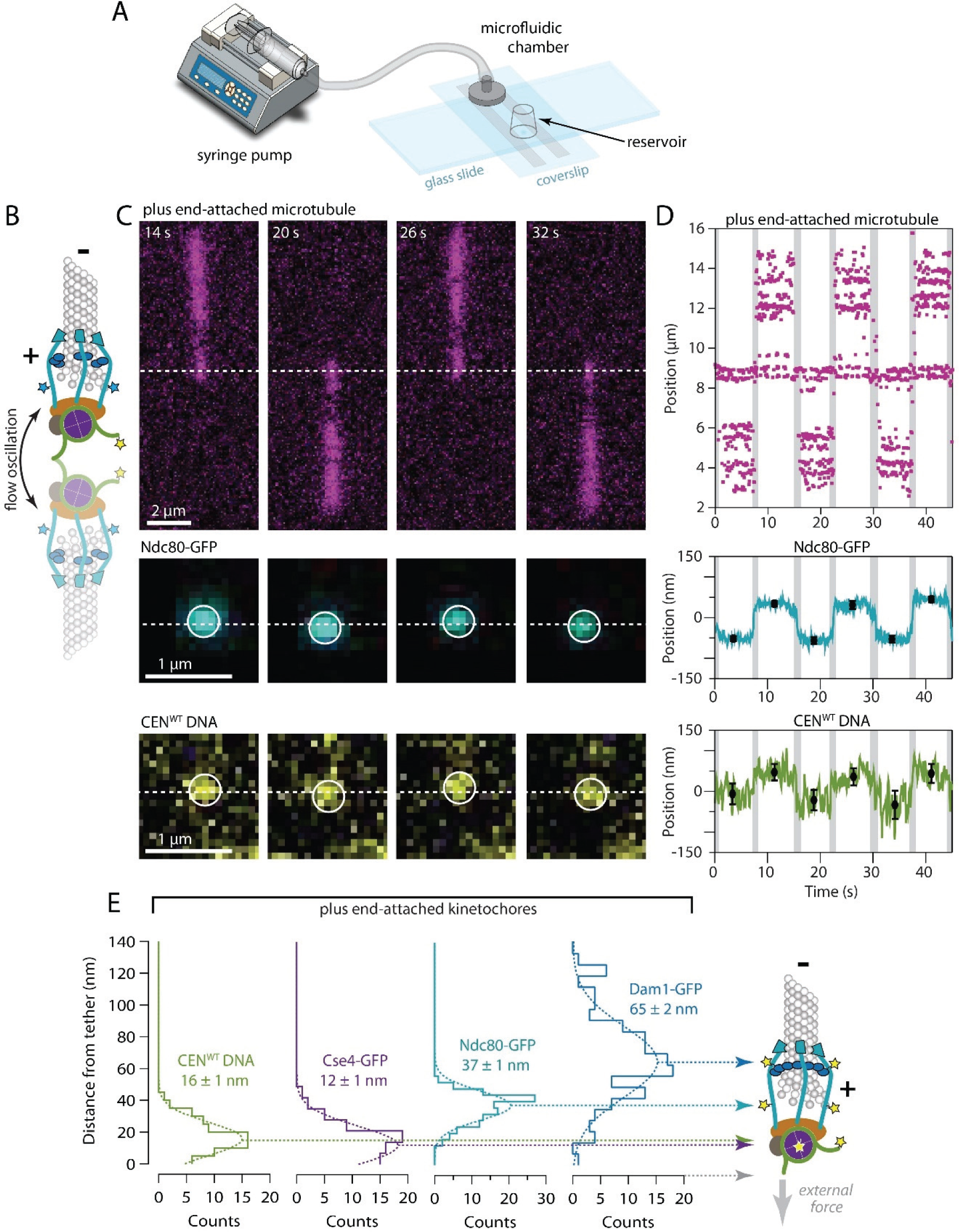
Plus end-attached kinetochores are well organized along the microtubule axis. **A)** Kinetochores were assembled in a microfluidic device and then allowed to capture microtubules. A syringe pump enabled imaging of the kinetochores and their captured microtubules while buffer was flowed gently through the assembly chamber. **B)** Schematic of a surface-assembled kinetochore attached to the tip of a microtubule. Oscillating the direction of flow caused the kinetochore and its captured microtubule to flip back and forth, reorienting by 180° with each reversal of the flow. **C)** Time-lapse image series showing flow-induced reorientation of a microtubule (magenta) attached by its end to a surface-assembled kinetochore. Both the Ndc80-GFP kinetochore marker (cyan) and the Atto565 label on the wild type centromeric DNA (yellow, CEN^WT^) oscillated with the direction of buffer flow. **D)** Example records of position versus time for an Ndc80-GFP spot and the corresponding Atto565-labeled centromeric DNA obtained by tracking the individual spots with sub-pixel accuracy. Displacements of each spot from the tether point were estimated by averaging during the intervals when the microtubule orientation was steady. Black symbols represent mean ± SDEV from *N* ≈ 60 tracked positions during each interval. Positions recorded during reorientation of the microtubule were omitted from the averaging and are indicated here by gray shading. Additional records are shown in Supplementary Figure 3s1. **E)** Distributions of displacement for the indicated fluorescent kinetochore components (from *N* = 67 to 128 intervals), fit with single Gaussian functions. The mean ± SEM for each Gaussian is indicated. Displacements for Cse4-GFP, a component of the centromeric nucleosome, are similar to the centromeric DNA (CEN^WT^), as expected. The larger displacements for outer microtubule-binding components, Ndc80-GFP and Dam1-GFP, are consistent with the *in vivo* arrangement.^58,65^

When a kinetochore periodically reoriented with the flow, the fluorescent marker on its centromeric DNA was displaced from the tether point on the coverslip by 16 ± 1 nm on average (Figures 3D and 3E; mean ± SEM from *N* = 74 measurement intervals across 13 end-attached kinetochores). This distance is much shorter than would be expected for a straight B-form DNA helix of 180 bp (∼61 nm), presumably because the centromeric DNA, after kinetochore assembly, was tightly wrapped around a centromeric nucleosome. Consistent with this interpretation, the histone H3 variant Cse4-GFP was located 12 ± 1 nm from the tether (mean ± SEM, *N* = 67 intervals, 9 kinetochores), very close to the centromeric DNA marker. The microtubule-binding component Ndc80-GFP was 37 ± 1 nm from the tether (*N* = 116 intervals, 19 kinetochores), implying that its C-terminal GFP tag was located ∼25 nm outward from the nucleosome (Figures 3D and 3E). Considering where the GFP tag falls within the structure of the Ndc80 complex,^61–63^ this 25-nm distance suggests the Ndc80c fibrils were well aligned with the microtubule axis. Dam1-GFP, a component of the outer, microtubule-binding Dam1 complex, was located 65 ± 2 nm from the tether (*N* = 128 intervals, 22 kinetochores) (Figures 3E and S5). This relatively large distance further suggests axial alignment of Ndc80c fibrils because the Dam1 complex binds nearer to the N-terminus of Ndc80.^33,64^ The implied intra-kinetochore separation between the C-termini of Ndc80 and Dam1 was 28 nm, a distance indistinguishable from the intra-kinetochore separation measured previously in budding yeast during metaphase.^58^ Altogether, these observations show that when *de novo* assembled kinetochores are attached to microtubule plus ends, they are spatially organized in a configuration that closely resembles the molecular arrangement of plus end-attached kinetochores during metaphase *in vivo*, with DNA-binding subcomplexes proximal to the chromatin and microtubule-binding subcomplexes projecting distally, toward minus ends.^58,65,66^

### Kinetochore architecture is less organized when attached to the sides of microtubules

The molecular organization of side-attached kinetochores has scarcely been explored. By mapping the relative positions of centromeric DNA and Ndc80-GFP within kinetochores that captured microtubules by their sides, we were able to examine the molecular arrangement of side-attached kinetochores and compare them directly to end-attached kinetochores, often measured simultaneously on the same coverslips. We focused on kinetochores that captured the sides of taxol-stabilized microtubules in an off-center arrangement (Figures 4A and 4B), where the two microtubule segments extending away from the kinetochore had unequal lengths. The longer segment experienced higher viscous drag forces and therefore oriented reliably downstream in the flow (Video S2). Axial positions of fluorescent-tagged centromeric DNA and Ndc80-GFP within these side-attached kinetochores were tracked in the same manner as for end-attached kinetochores (Figure 4C). The distribution of distances between the centromeric DNA marker and the tether point on the coverslip was indistinguishable from that measured for end-attached kinetochores (Figures 4D and 4E; *p* = 0.15, based on a K-S test with *N* = 32 and 74 intervals from 13 side- and 13 end-attached kinetochores, respectively). The distribution of Ndc80-GFP distances, however, was wider in comparison to the end-attached kinetochores and apparently bimodal, including an elongated sub-population with a mean distance of 39 ± 4 nm and a compact sub-population, much closer to the tether, with a mean distance of only 18 ± 9 nm (± SEM, *N* = 54 and 33 intervals, respectively, 13 side-attached kinetochores) (Figure 4E). This observation shows that the molecular architecture of side-attached kinetochores differs from end-attached kinetochores, with Ndc80 fibrils often less well aligned to the microtubule axis.

**Figure 4.**
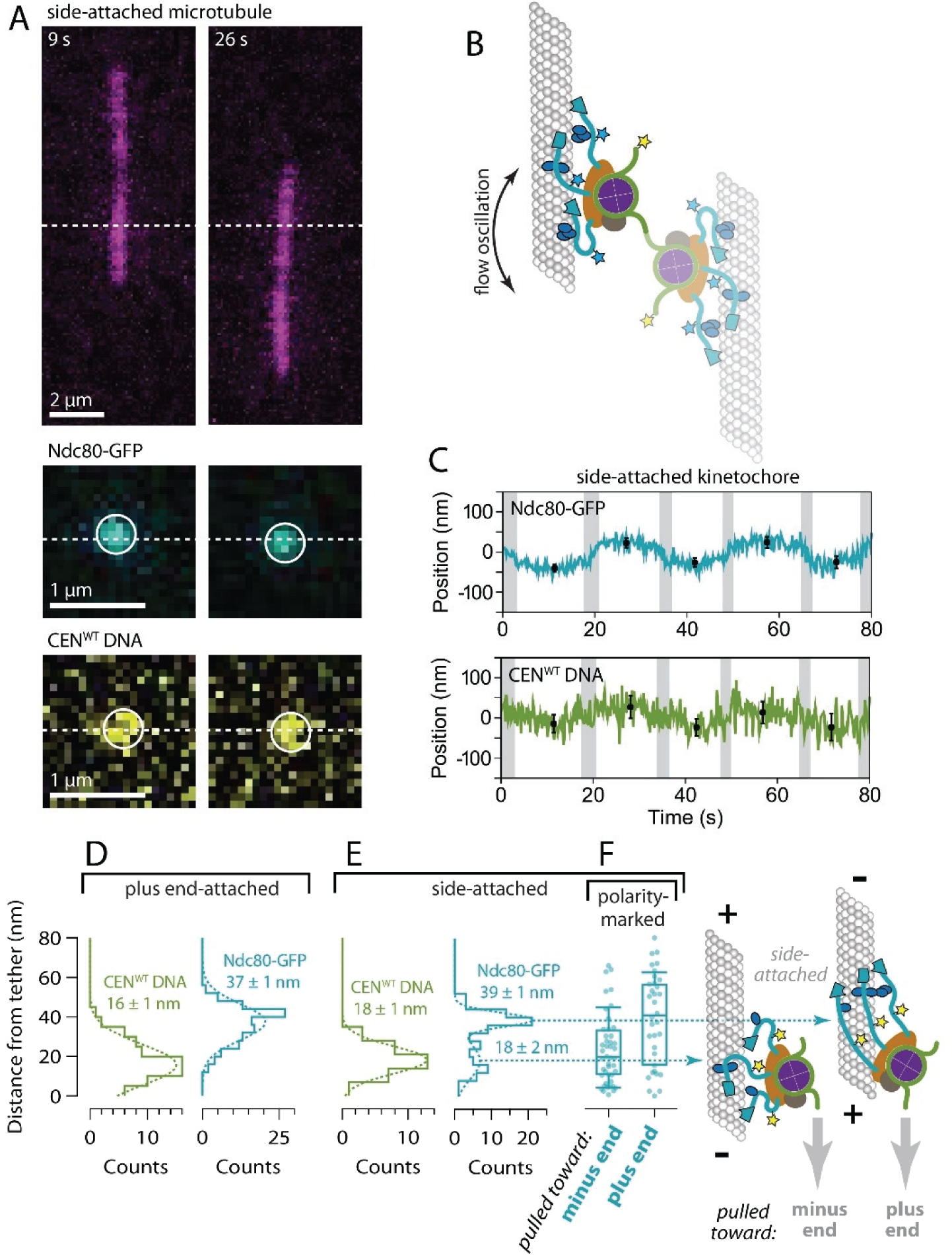
Side-attached kinetochores are more compact specifically when pulled toward minus ends. **A)** Time-lapse images showing flow-induced reorientation of a microtubule (magenta) attached by its side to a surface-assembled kinetochore. Both the Ndc80-GFP kinetochore marker (cyan) and the Atto565 label on the centromeric DNA (yellow) oscillated with the direction of buffer flow. **B)** Schematic of a surface-assembled kinetochore attached to the side of a microtubule. Oscillating the direction of flow caused the kinetochore and its captured microtubule to flip back and forth, reorienting by 180° with each reversal of the flow. **C)** Example records of position versus time for an Ndc80-GFP kinetochore attached to the side of a microtubule. Displacements of both the Ndc80-GFP spot and the Atto565 label on the wild type centromeric DNA (CEN^WT^) relative to the tether point were estimated by averaging during the intervals when the microtubule orientation was steady. Black symbols represent mean ± SDEV from *N* ≈ 60 tracked positions during each interval. Positions recorded during reorientation of the microtubule were omitted from the averaging and are indicated here by gray shading. **D)** Distributions of displacement for the indicated fluorescent components within tip-attached kinetochores (from *N* = 74 to 116 intervals), fit with single Gaussian functions. The mean ± SEM for each Gaussian is indicated. These data and fits are replotted from Figure 4B with an expanded vertical scale. **E)** Distributions of displacement for the indicated fluorescent components within side-attached kinetochores (from *N* = 32 to 87 intervals), fit with either a single Gaussian (CEN^WT^ DNA) or a double Gaussian function (Ndc80-GFP). The mean ± SEM for each Gaussian is indicated. The distribution of Ndc80-GFP displacements for side-attached kinetochores is wider in comparison to the tip-attached kinetochores in panel D, and apparently bimodal, including a sub-population very close to the tether with a mean displacement of only 18 ± 2 nm. **F)** Distributions of displacement for Ndc80-GFP within kinetochores attached to the sides of polarity-marked microtubules. Boxes extend from the first to the third quartiles with medians indicated by the central horizontal solid lines. Medians for kinetochores pulled toward plus and minus ends were 40.9 and 19.6 nm, respectively (from *N* = 34 and 46 intervals). Whiskers extend ± one SDEV from the mean.

### Side-attached kinetochores are more compact specifically when pulled toward minus ends

Depending on which end of the microtubule was oriented downstream in the flow, the side-attached kinetochores experienced sub-piconewton viscous pulling forces directed either toward the minus end or toward the plus end. We hypothesized that the two sub-populations, with elongated or compact Ndc80-GFP, might correspond to these two different pulling directions. To test this idea we repeated the Ndc80-GFP distance measurements using polarity-marked GMPcPP-stabilized microtubules. When side-attached kinetochores were pulled toward minus ends Ndc80-GFP was closer to the tether, at a distance of only 24 ± 3 nm (mean ± SEM, *N* = 46 intervals, 12 kinetochores), and when they were pulled toward plus ends Ndc80-GFP was farther from the tether, at a distance of 39 ± 4 nm (*N* = 34 intervals, 7 kinetochores; *p* = 0.003 based on a K-S test) (Figure 4F). These measurements demonstrate that the direction of external force directly influences kinetochore architecture, with Ndc80 fibrils adopting a more compact arrangement specifically when the kinetochore is pulled toward the minus end.

## DISCUSSION

Given the vital importance of plus-end attachments for chromosome segregation, an attractive idea has been that kinetochores possess an intrinsic preference for plus ends over other regions of the microtubule. Various purified kinetochore subcomponents bind preferentially to curved tubulin^40,67,68^ or GTP caps.^31^ However, whether these behaviors confer a plus end preference to the kinetochore as whole is unclear. Our isolated kinetochores captured microtubules with a strong preference for plus ends and their attachment strength was higher at plus ends than at minus ends. We can exclude a role for active motors in these behaviors because our kinetochores lacked motors^14,53^ and ATP was absent from our experiments. Our capture assays used filaments stabilized by GMPcPP and taxol, indicating that preferential plus end capture does not require GTP caps or microtubule dynamics, and probably does not require curved tubulin since the ends of GMPcPP microtubules are usually blunt.^69^ Moreover, we found that side-attached kinetochores were sensitive to microtubule polarity even far away from the ends of the filaments. We propose that all these intrinsic kinetochore behaviors – their preference for capturing and holding microtubule plus ends, and their directionally asymmetric grip when side-attached – arise from the structural polarity of the microtubule and how it influences kinetochore architecture (Figure 5).

**Figure 5.**
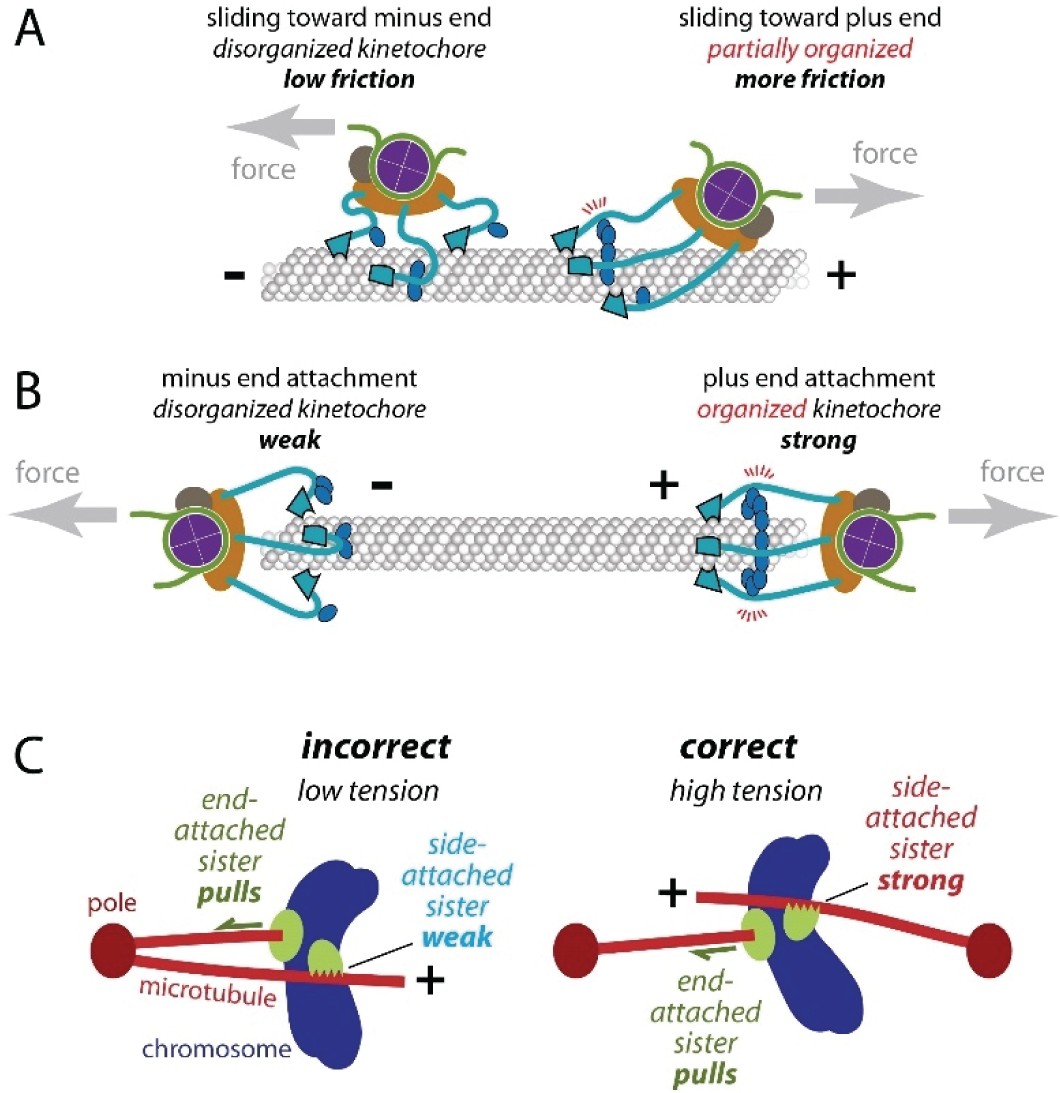
How the directionally asymmetric grip of the kinetochore might arise from microtubule polarity and promote accuracy during early mitosis. **A)** Each Ndc80c fibril (light blue) has a globular ‘foot’ (outlined in black), which binds with a stereospecific ‘footprint’ on the outside surface of the microtubule, and a coiled-coil stalk that emanates from this foot and projects toward the plus end. *Right:* Pulling a kinetochore toward the plus end aligns its multiple Ndc80c stalks into a parallel configuration, facilitating interactions with Dam1c oligomers and strengthening the overall grip on the microtubule. Because the direction of force is aligned with the stalks, torque on the Ndc80c-microtubule bonds is minimized. *Left:* Pulling a kinetochore toward the minus end disrupts this organization, weakening its grip. **B)** *Right:* At a plus end, the Ndc80c stalks can all project in parallel past the tip of the microtubule and converge onto the centromeric nucleosome,^35^ potentially allowing Dam1c oligomers to organize a cage-like arrangement surrounding the tip,^32,35^ with even higher grip strength. *Left:* At a minus end, parallel convergence of the stalks is impossible, increasing the torque on the Ndc80c-microtubule bonds and potentially reducing lateral interactions via Dam1c. **C)** After a pair of sister kinetochores initially makes side-attachments, one of them will (by chance) become tip-attached before the other, tracking with tip shortening and exerting elastic pulling forces on its side-attached sister. *Left:* If the pair is attached incorrectly to microtubules from the same pole, then the side-attached sister will be pulled toward the minus end. Its grip will therefore be weak, and it will likely detach. *Right:* If the pair is attached correctly to microtubules from opposite poles, then the side-attached sister will be pulled toward the plus end. It will therefore have a stronger grip that should allow it to remain attached and achieve proper bi-orientation at the plus end.

Ndc80c binds microtubules partly through a stereospecific ’footprint’, which presumably cannot twist or rotate without breaking its bond to the microtubule. The stalk of Ndc80c emerges from the ‘foot’ (from the calponin-homology domains) with a tilt toward the microtubule plus end.^28,35,70^ This local structural asymmetry probably sets up the asymmetric mechanical behavior of the whole kinetochore. When a side-attached kinetochore is pulled toward the plus end, its Ndc80c feet may bind more strongly. When pulled toward the minus end, its feet may bind more weakly. In any case, after emerging asymmetrically from the foot, the Ndc80c stalk contains a flexible ‘hinge’,^63,71,72^ which suggests that the rest of the stalk should align at least partially with the direction of external force. Indeed, our data imply that pulling a kinetochore toward the plus end aligns its Ndc80c stalks into a parallel configuration, potentially facilitating interactions with Dam1c oligomers that can provide additional microtubule contacts to strengthen the overall grip of the kinetochore on the microtubule (Figure 5A).[refs?] At a plus end, the stalks should all project past the tip of the microtubule to converge onto the centromeric nucleosome,^35^ potentially allowing Dam1c oligomers to organize a cage-like arrangement surrounding the tip,^34,35^ and further increasing grip strength (Figure 5B). Pulling a kinetochore toward the minus end disrupts the parallel organization of Ndc80c stalks, likely preventing Dam1c from making additional microtubule contacts, weakening the kinetochore’s grip when side-attached, and also precluding strong attachment to the minus end.

Independent of the molecular underpinnings, the directionally asymmetric grip of the kinetochore suggests a previously unrecognized mechanism for promoting accuracy early in mitosis. The accuracy of mitosis is astounding,^73,74^ and relies on a trial-and-error process where incorrect attachments lacking tension are released while correct tension-bearing attachments are stabilized.^8–10^ The mechanisms proposed to explain this selectivity have focused primarily on plus end-attachments.^9–11^ However, kinetochores first bind the sides of spindle microtubules,^1–5^ a behavior that facilitates microtubule capture because a much greater surface area is available along the sides than at the ends of the microtubules. Whether erroneous side-attachments are distinguishable from correct side-attachments during this early attachment phase has scarcely been considered. Our findings now indicate that the discrimination between correct and incorrect attachment can begin even before both sisters have achieved plus end-attachments. In prometaphase, sister kinetochores are exposed to many spindle microtubules emanating from both poles. After a pair of sister kinetochores initially makes side-attachments, one of them will (by chance) become tip-attached before the other, tracking with tip shortening and exerting elastic pulling forces on its side-attached sister. If the pair is attached *incorrectly* to microtubules from the same pole, then the side-attached sister will be pulled toward the minus end. Its grip will therefore be weak, and it will likely detach. Conversely, if the pair is attached *correctly* to microtubules from opposite poles, then the side-attached sister will be pulled toward the plus end. It will therefore have a stronger grip that should allow it to remain attached and achieve proper bi-orientation at the plus end. The greater frictional resistance of the correctly side-attached sister will also cause the end-attached kinetochore to experience higher force, stabilizing its attachment by the catch bond-like effect we previously discovered,^14^ and protecting it from Aurora B kinase-triggered detachment.^13^ Thus the asymmetric grip of the side-attached sister can selectively stabilize correct over incorrect arrangements, prior to plus-end bi-orientation, in several ways.

We note also that the directionally asymmetric grip of the kinetochore is similar to the recently discovered directional binding of focal adhesion proteins to F-actin.^41–43^ This similarity suggests that asymmetric gripping may be a general mechanism for selective attachment of cytoskeletal filaments with appropriate orientation at cytoskeletal junctions in many cellular contexts.

## ACKNOWLEDGEMENTS

We thank Trisha Davis, Bonnibelle Leeds, Lucia Kyung-Ae Maki-Fern, and Juan-Jesus Vicente for critical reading and feedback on the manuscript, and all the members of the Davis, Asbury, and Biggins labs for insightful discussions. We also thank Christian Nelson and Cameron Lee for constructing strains and purifying kinetochores. This project was funded by the National institutes of Health (T32HL007312 to J.D.L., F32GM136010 to A.R.P., R01GM079373 and R35GM134842 to C.L.A., and R01GM064386 to S.B.), a Washington Research Foundation Postdoctoral Fellowship to J.D.L., and a Mary Gates Research Scholarship to N.A.H. S.B. is an investigator of the Howard Hughes Medical Institute.

## AUTHOR CONTRIBUTIONS

Conceptualization, J.D.L., S.B., and C.L.A.; Methodology, Investigation, and Formal Analysis, J.D.L., N.A.H., L.E.M. C.L.A.; Software, C.L.A.; Recourses, J.D.L., A.R.P., S.B.; Writing – Original Draft, J.D.L, L.E.M., and C.L.A.; Writing – Review & Editing, J.D.L, N.A.H., L.E.M., A.R.P., S.B., and C.L.A.; Funding Acquisition, J.D.L., N.A.H. and C.L.A.; Supervision, J.D.L., S.B. and C.L.A.

## MATERIALS AND METHODS

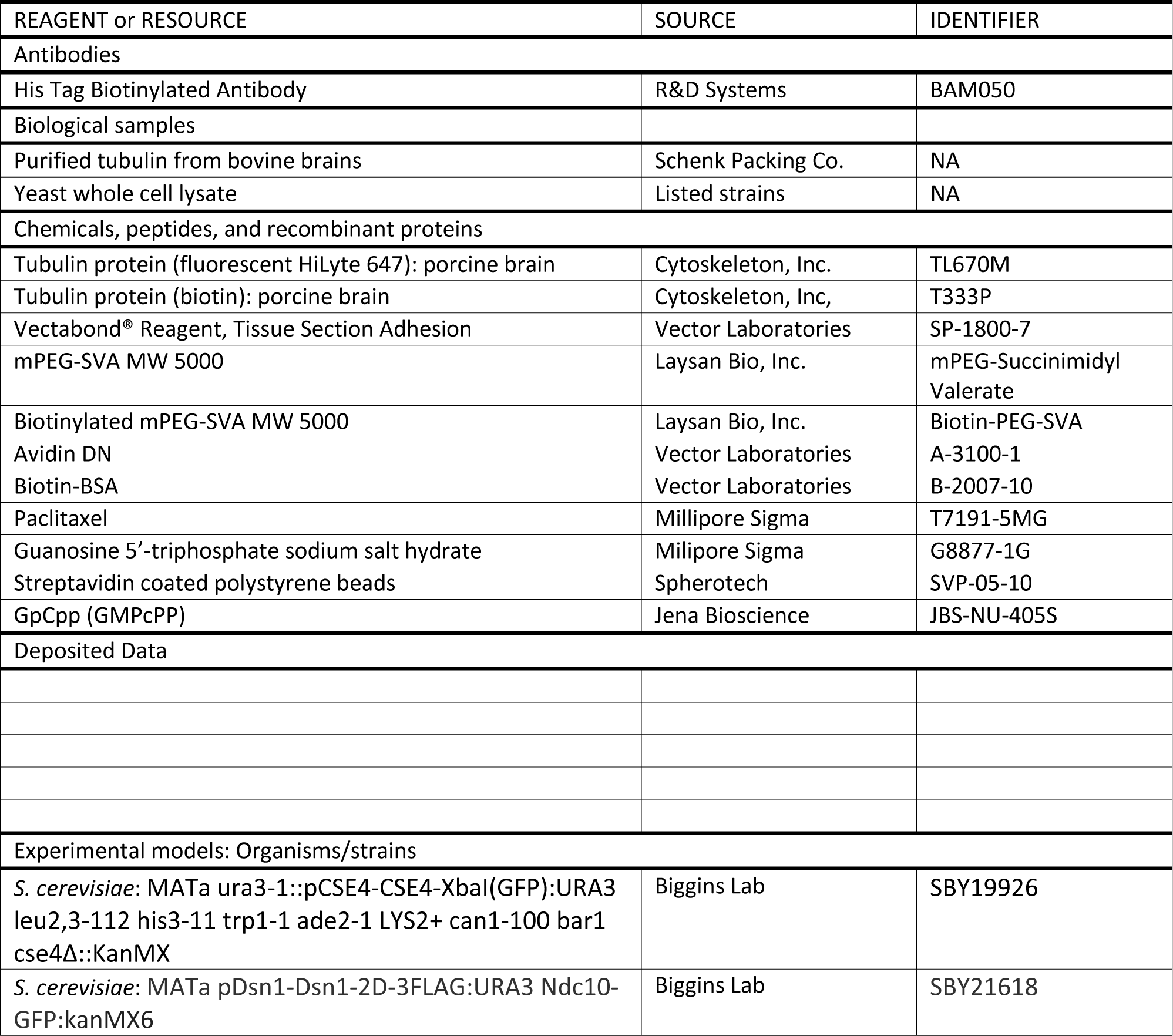
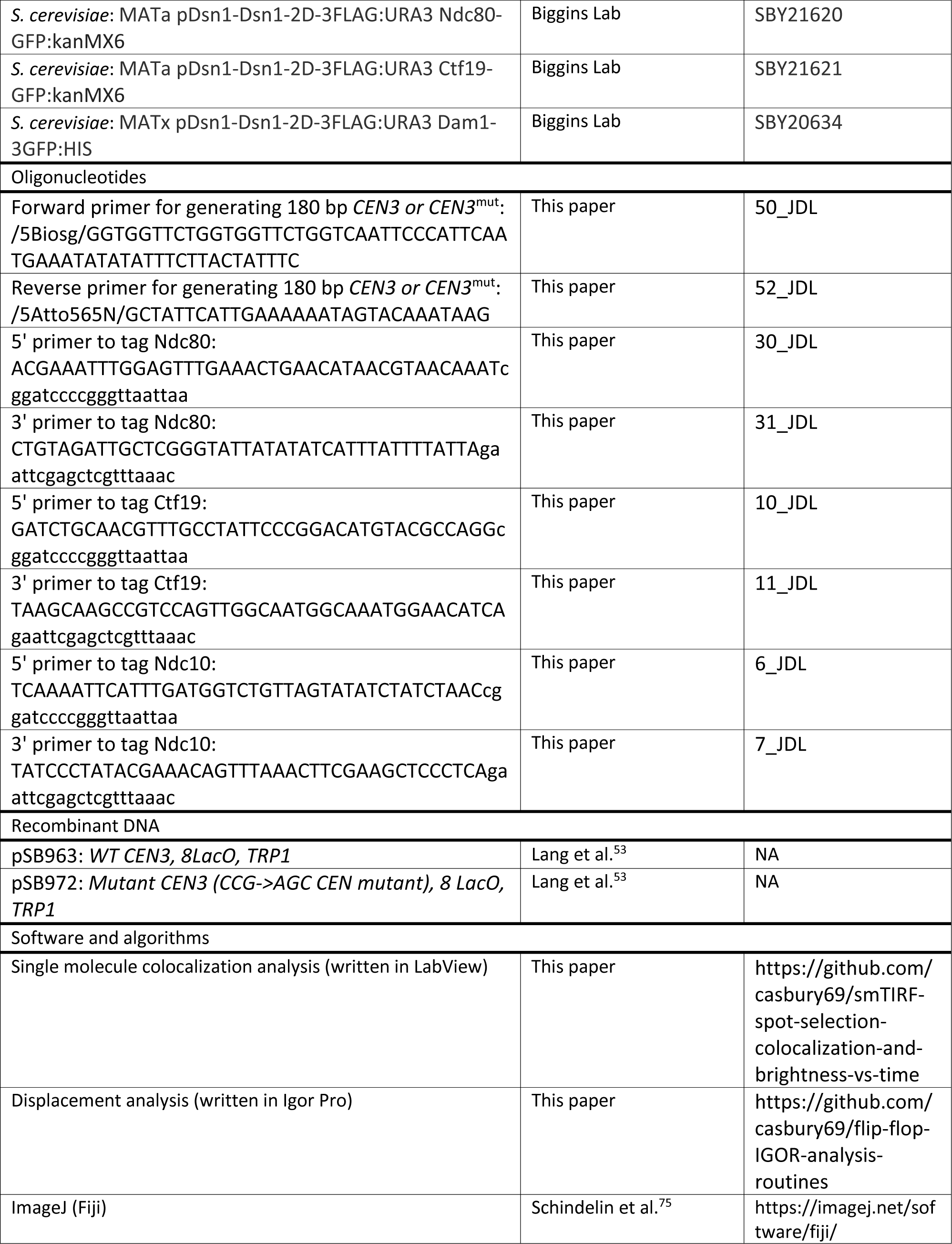

### Yeast strain construction

All strains described in this study are derivatives of SBY3 (W303). Generation of *Saccharomyces cerevisiae* strains harboring GFP tagged kinetochore proteins and a phospho-mimetic mutation in Dsn1 (Dsn1-2D) that has been shown to enhance outer kinetochore assembly^53,77^ was achieved either by standard genetic crosses and media selection^78^ or using standard PCR-based integration at the endogenous loci.^79^ All yeast transformants were confirmed by PCR or sequencing.

### Preparation of yeast whole cell lysates

Yeast whole cell lysates used for *de novo* kinetochore assembly were prepared as previously described.^46,53^ Cells were grown in 2 L of liquid yeast peptone dextrose (YPD) media at room temperature and harvested in log phase by centrifugation. Cell pellets were placed on ice and washed with ice cold dH_2_0 plus 0.2 mM PMSF and centrifuged. Pellets were washed a second time with ice cold Buffer L (25 mM HEPES pH 7.6, 2 mM MgCl2, 0.1 mM EDTA, 0.5 mM EGTA, 0.1% NP-40, 175 mM K-Glutamate, and 15% Glycerol) plus 2 mM DTT and protease inhibitors (10 µg/mL leupeptin, 10 µg/mL pepstatin, 10 µg/mL chymostatin, and 10 µM PMSF) and centrifuged again. Cells were resuspended in a final volume of Buffer L given by *u* = *v* · *o*, where *u* represents the volume in µL of added Buffer L, *v* represents the original volume in mL of the liquid culture, and *o* represents the optical density of the culture measured at the time of harvest. The cellular resuspension was snap frozen as small spherical pellets by pipetting drops of the suspension directly into liquid nitrogen. Cell lysis was achieved using a Freezer/Mill (SPEX SamplePrep) by alternating milling of the pellets at 10 Hz for 2 minutes followed by a 2-min cooling phase for ten cycles. The resulting lysate powder was then thawed on ice and clarified by centrifugation at 16,100 g for 30 minutes at 4°C. The protein-containing supernatant was subsequently aliquoted (100 µL) and snap frozen in liquid nitrogen. Aliquots were stored at -80°C until use.

### Preparation of CEN DNAs

180 base pair Atto565 CEN^WT^ and CEN^mut^ DNA were generated by PCR from plasmids pSB963 and pSB972, respectively. CEN^mut^ DNA contained a three base pair substitution in the CDEIII region of CEN3 that blocks kinetochore assembly in vivo^50–52^ and *in vitro*.^53^ The forward 5’ primer contained a 5’ biotin for attachment to the PEG/biotin-PEG passivated coverslip and the reverse primer was labeled with Atto565. PCR products were purified using a Qiagen PCR Cleanup kit and eluted into dH_2_0.

### Preparation of taxol-stabilized microtubules

Purified bovine brain tubulin was added to microtubule polymerization buffer [1x BRB80 (80 mM PIPES, 1 mM MgCl2, 1 mM EGTA), 7% DMSO, 4 mM MgCl_2_, and 1 mM GTP] to a final concentration of 2 mg/mL and incubated at 37°C for 1 hr. After 1 hour, 3 µL of pre-warmed 1x BRB80 + 10 µM taxol was added for every 1 µL of polymerized microtubules. Taxol-stabilized microtubules were then spun for 10 min at 90,000 RPM (TLA100.0, Beckman Optima MAX-XP) at 37°C. The microtubule pellet was resuspended in 150 µL of 1x BRB80 + 10 µM taxol and stored at room temperature. To generate fluorescent or biotinylated microtubules, porcine HyLite 647, or biotin tubulin (Cytoskeleton) were added (6% w/w) to the polymerization reaction.

### Slide passivation for single molecule TIRF microscopy

Slides were prepared as previously described by Crawford et al.^80^ Coverslips and slides were plasma cleaned for 10 min followed by four sequential hour-long sonications in 2% Micro-90, 200 proof ethanol, 1 M KOH, and finally Milli-Q water. Slides and coverslips were then completely dried using ultrapure nitrogen. After drying, slides and coverslips were treated with Vectabond (Vector Laboratories) dissolved in acetone (1% v/v) for 5-10 min. Slide chambers were constructed by sandwiching the coverslip and slide together with double-sided tape. Passivation was achieved by adding a 1:100 Biotinylated mPEG-SVA/mPEG-SVA MW 5000 in 0.1 M sodium bicarbonate (1% w/v). Chambers were incubated with PEG overnight at room temperature.

### Kinetochore assembly and microtubule capture assays

Excess mPEG solution was washed out with 400 µL of 1x BRB80 and then blocked with a 0.1 mg/mL BSA solution for 5 min. The chamber was then washed with 200 µL of 1x BRB80. After blocking, a solution of 0.3 mg/mL avidin was added to the chamber for 5 min and washed with an additional 200 uL of 1x BRB80. Following the addition of avidin, 50-200 pM biotinylated Atto565 CEN DNAs were introduced into the chamber and incubated for 5 min. Excess DNA was washed away with 200 µL of 1x BRB80. To assemble surface tethered kinetochores, 100 µl of yeast whole cell lysate was added to chambers with surface tethered CEN DNAs and incubated for 1 hr. For colocalization assays, the lysate was washed away with 400 µL of 1x BRB80 with glucose oxidase (165 U/mL)/catalase (217 U/mL) and 0.65% glucose (w/v) for scavenging oxygen. For microtubule pulldown assays, taxol-stabilized microtubules were diluted 1:3 in BRB80 + 10 µM taxol and briefly sheared using a vortexer for 25 s before introduction to the slide chamber. After a 15-min incubation, excess microtubules were washed away with 400 µL of 1x BRB80 with glucose oxidase/catalase.

### Single-molecule colocalization analysis

All images were collected on a custom TIRF microscope built on a standard Nikon TE inverted microscope base.^81^ Excitation of fluorescent proteins and organic dyes was achieved using expanded beams from three solid-state lasers at 488 nm (Coherent Sapphire), 561 nm (Coherent Sapphire) and 641 nm (Coherent Cube). Images were acquired with three separate Andor iXon897+ EMCCD cameras. For colocalization assays, 20 to 60 frames were collected with 0.5 second integrations. Analysis was performed using custom Labview (National Instruments) software available at https://github.com/casbury69/smTIRF-spot-selection-colocalization-and-brightness-vs-time. The software implements spot picking for each fluorescent channel using methods described by Crocker and Grier,^82^ and by Friedman and Gelles.^83^ Mapping between color channels was performed by creating a linear registration map using blue/green/orange/dark red 500 nm beads (Tetraspec, T7281) as fiducials.

### Preparation of polarity marked microtubules

To produce polarity marked microtubules, two seed mixes were prepared on ice: a bright seed mix (13.3 µM unlabeled bovine tubulin, 6.7 µM Hilyte 647 tubulin, 1 mM DTT, 1 mM GMPcPP, 1x BRB80) and a dim seed mix (9 µM unlabeled bovine tubulin, 1 µM HyLite 647 cytoskeletal tubulin, 8 µM NEM-treated bovine tubulin, 1 mM DTT, 1 mM GMPcPP, 1x BRB80). Each seed mix was clarified using an ultra-centrifuge (Beckman Optima MAX-XP, 90,000 RPM, 4°C, 5 min) then snap-frozen in small aliquots and stored at -80°C. Bright seeds were polymerized by diluting an aliquot of bright seed mix 5-fold (vol/vol) in 1x BRB80 with 1 mM DTT. Diluted bright seed mix was then incubated for 45 min at 37°C to allow for polymerization. To grow dim elongations from the bright seeds, an aliquot of dim seed mix was diluted 5.7-fold (vol/vol) in 1x BRB80 with 1 mM DTT on ice, then warmed for 20 s at 37°C. Polymerized bright seeds were added 4.4-fold (vol/vol) to the dim mix and incubated for 1 hr at 37°C. After the second polymerization with the dim seed mix, the microtubules were centrifuged for 5 min at 22,000 RPM (TLA100.0, Beckman Optima MAX-XP) at 37°C. The pellet was then resuspended in 150 µL assembly assay buffer (1xBRB80, 1 mM DTT, 0.025 mg/mL K-casein, 20 µM taxol).

### Tracking kinetochore subunit displacements and estimating intra-kinetochore distances

Custom flow chambers, with an attached reservoir to hold excess buffer and with custom-made fittings, were designed to generate a gentle oscillating flow to orient surface assembled kinetochore/microtubule pairs along the coverslip surface. The fitting was attached to a syringe pump (Kd Scientific, 780210) which operated at a slow flow rate of 0.6 mL/min. The volume of each oscillation was 0.2 mL. Images were acquired in the same manner used during the colocalization assays with the exception of using 0.2 second integrations instead of 0.5 second integrations. The total number of frames collected was determined by the bleach rate of GFP for the individual kinetochores. Particle tracking was performed using the MOSAICsuite 2D particle tracker plugin for ImageJ.^76^ Analysis of particle displacements was achieved using custom scripts written in IgorPro (Wavemetrics) and available at https://github.com/casbury69/flip-flop-IGOR-analysis-routines.

### Rupture force assay

Rupture force assays were carried out as previously described.^14,15^ Dynamic microtubules were grown from biotinylated-GMPCPP seeds on biotinylated-BSA passivated coverslips in microtubule growth buffer (BRB80, 1 mM GTP, 250 µg/mL glucose oxidase, 25 mM glucose, 30 µg/mL catalase, 1 mM DTT, 24 µM purified bovine brain tubulin, 0.5 mg/mL κ-casein). Anti-HIS antibody (R&D systems, BAM050) coated polystyrene beads were prepared as previously described.^39^ The beads were decorated with native purified Dsn1-6HIS-3Flag kinetochores, and bound to dynamic plus or minus end microtubule tips. A feedback-controlled laser trap was used to apply increasing force to tip bound kinetochore decorated beads in the direction of microtubule assembly. The applied force was increased at a constant rate of 0.25 pN/s until the bead detached from the microtubule tip or the force exceeded the limit of the laser trap. Bead position was recorded using custom LabView software and analyzed to determine the rupture force with custom scripts in IgorPro. The longest extension from the GMPcPP seeds was designated as the plus end, and the shorter extension was designated as the minus end.

### Drag force assay

Microtubules were polymerized as described above with the addition of a ten-minute incubation with growth buffer containing 24 µM tubulin to allow for minus end extensions to grow enough for bead binding along the microtubule side. The laser trap was then used to apply a constant force in one direction along the longitudinal axis of the microtubule for a short distance before reversing the direction of the applied force. This was repeated several times at forces varying from 0.5 to 4 pN on both the minus and plus end extension of the microtubules. Velocity analysis was done using custom scripts in IgorPro.

## SUPPLEMENTAL FIGURE TITLES AND LEGENDS

**Figure S1.**
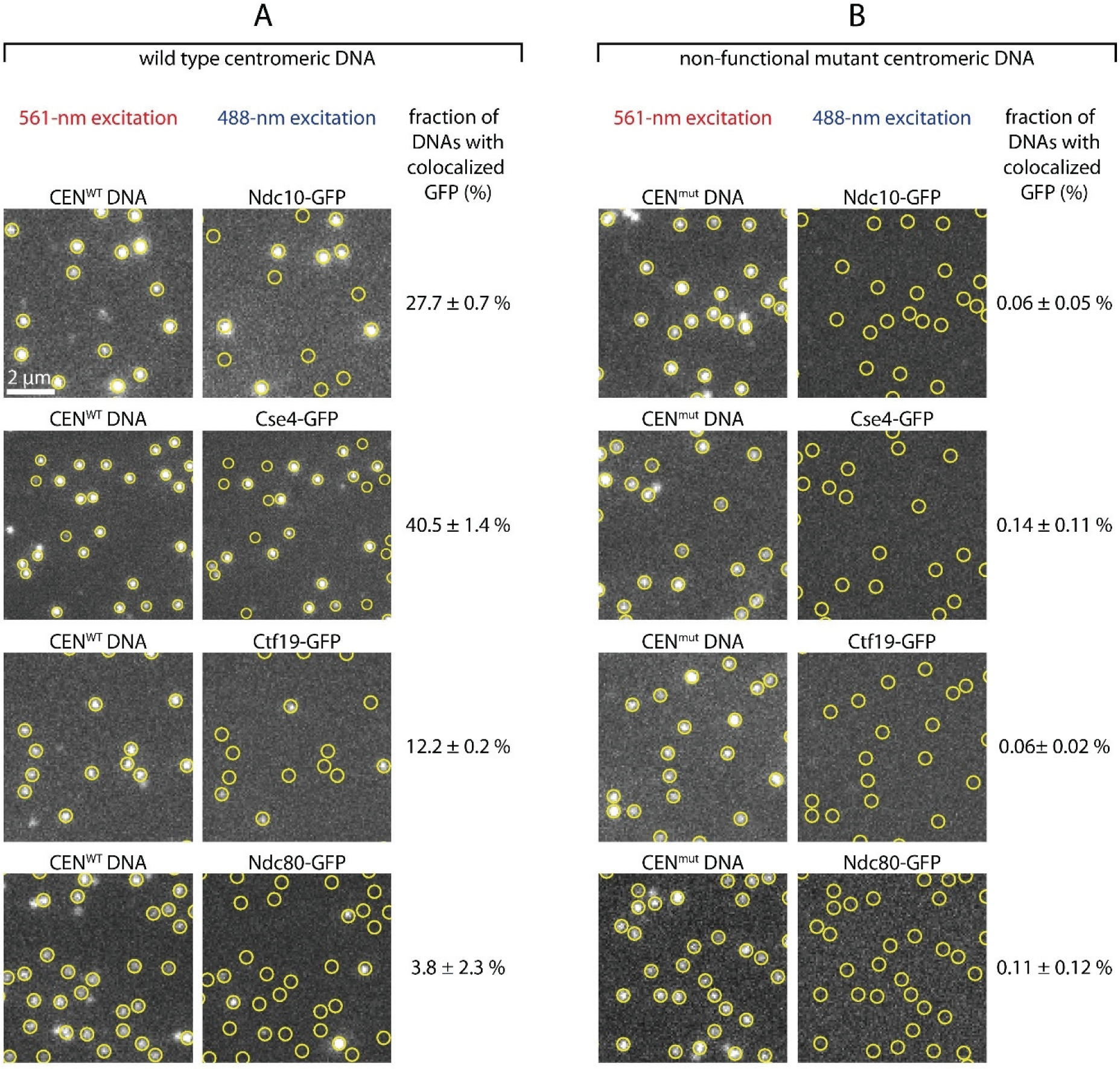
*De novo* assembly of individual kinetochores occurs specifically on centromeric DNAs. **A)** Images of individual Atto565-labeled centromeric DNAs (left) and corresponding images of GFP-tagged kinetochore subcomplexes (right) from the same fields of view. Yellow circles mark locations of individual wild type centromeric DNAs (CEN^WT^), which contain the complete 117-bp centromere sequence from *S. cerevisiae* chromosome III. **B)** Images from negative control experiments using mutant centromeric DNAs (CEN^mut^) carrying a 3-bp substitution that prevents kinetochore assembly. The percentages in both A and B represent average fractions (± SEM) of centromeric DNAs that colocalized with a GFP signal from the indicated kinetochore protein, calculated from *N* > 3,400 DNAs for each kinetochore component from at least 9 fields of view across three independent experiments.

**Figure S2.**
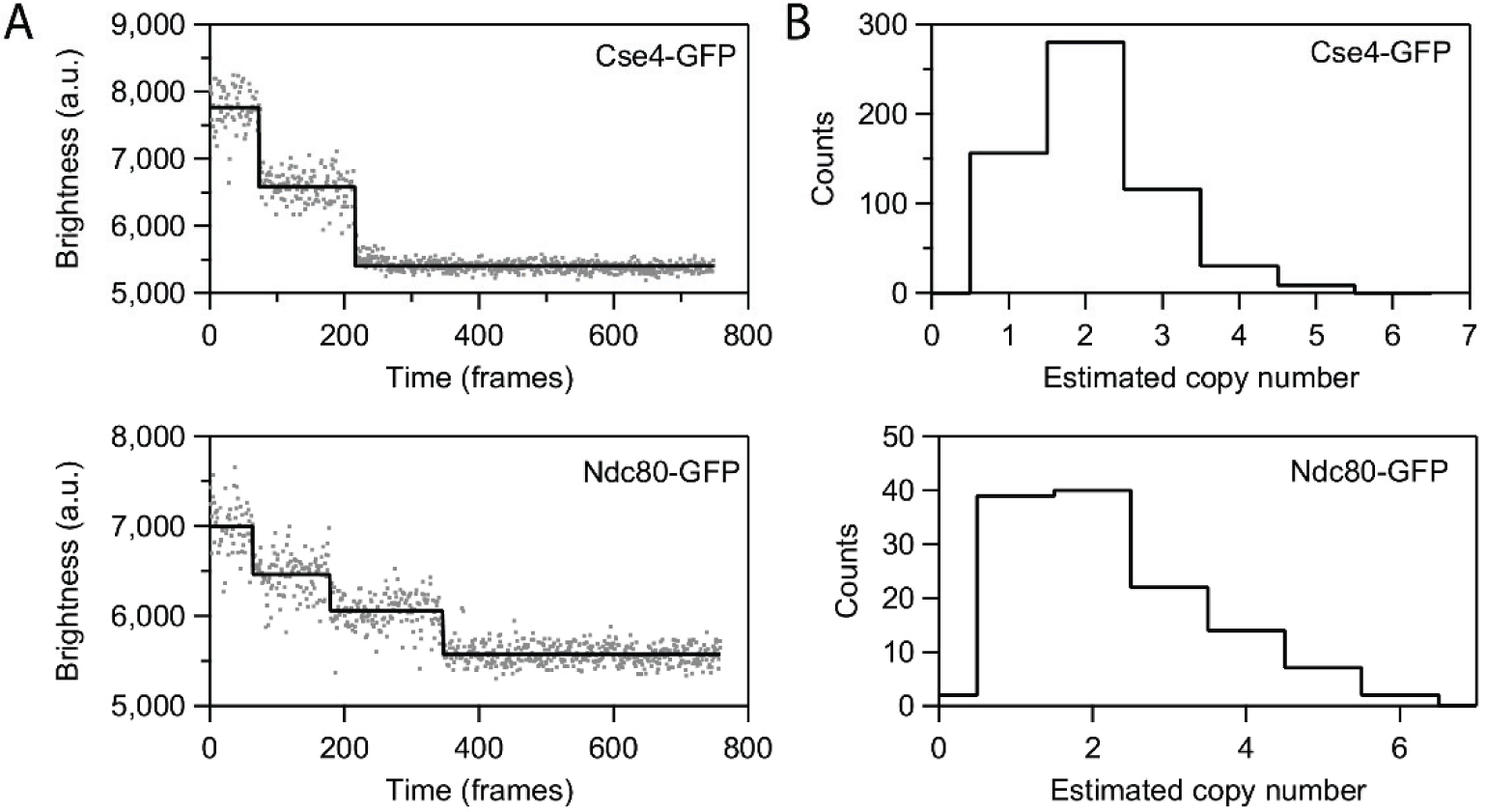
Photobleach analysis of Cse4- or Ndc80-GFP assembled kinetochores. **A)** Representative records of fluorescence intensity versus time for individual Cse4-GFP (top) or Ndc80-GFP (bottom) assembled kinetochores. The raw intensity data is represented by grey spots and the estimated bleach steps are represented by the solid black line. Bleach steps were estimated using the Tdetector2 step detection algorithm.^84^ **B)** Histograms showing the estimated copy number of Cse4-GFP (top) or Ndc80-GFP (bottom) present in individual assembled kinetochores. *N* = 591 individual Cse4-GFP kinetochores and *N* = 126 individual Ndc80-GFP kinetochores.

**Figure S3.**
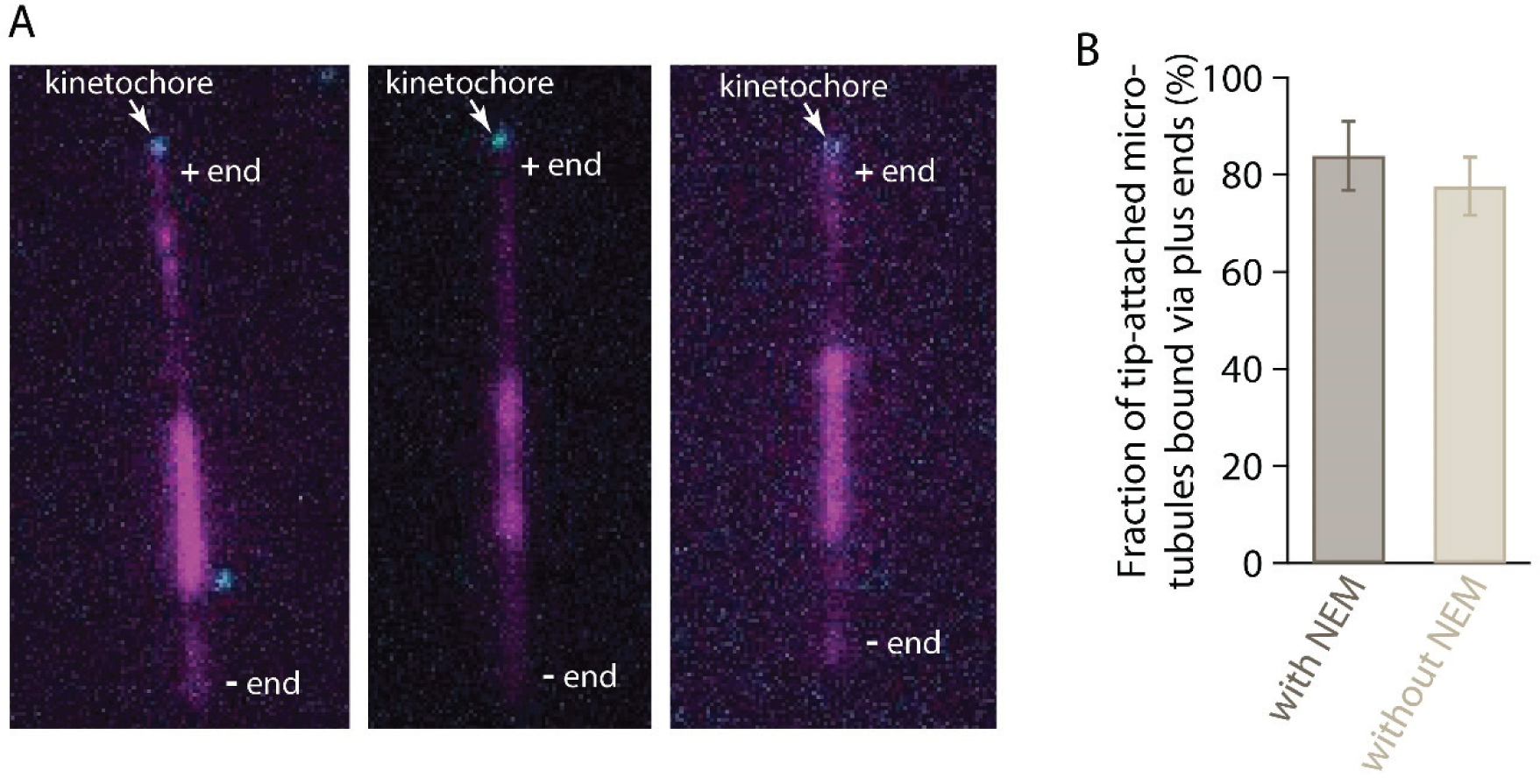
Plus end preference is not an artifact of differential labeling. **A)** Polarity-marked microtubules (magenta), polymerized from bright seeds with dim extensions on both plus and minus ends, were nearly always captured via their plus ends by assembled Ndc80-GFP kinetochores (cyan). To polymerize dim extensions from both ends of bright seeds, NEM-treated tubulin was omitted from the polymerization mix (see Materials and Methods). Polymerization at plus ends is faster than at minus ends, so plus ends were distinguishable by their longer lengths relative to minus ends. **B)** Percentages of tip-captured, polarity marked microtubules that bound by their plus ends. A strong preference for plus ends occurred irrespective of whether the minus ends were more brightly labeled, via polymerization with a small amount of NEM-treated tubulin (with NEM, at left), or whether the minus and plus ends were both dimly labeled, via polymerization without NEM-treated tubulin (without NEM, at right). See Materials and Methods for details about how polarity-marked microtubules were generated. Bars represent percentages of polarity-marked microtubules that were bound by their plus ends (mean ± SDEV, from *N* = 4 experiments with NEM and *N* = 4 without NEM, examining a total of 86 and 110 tip-captured microtubules, respectively).

**Figure S4.**
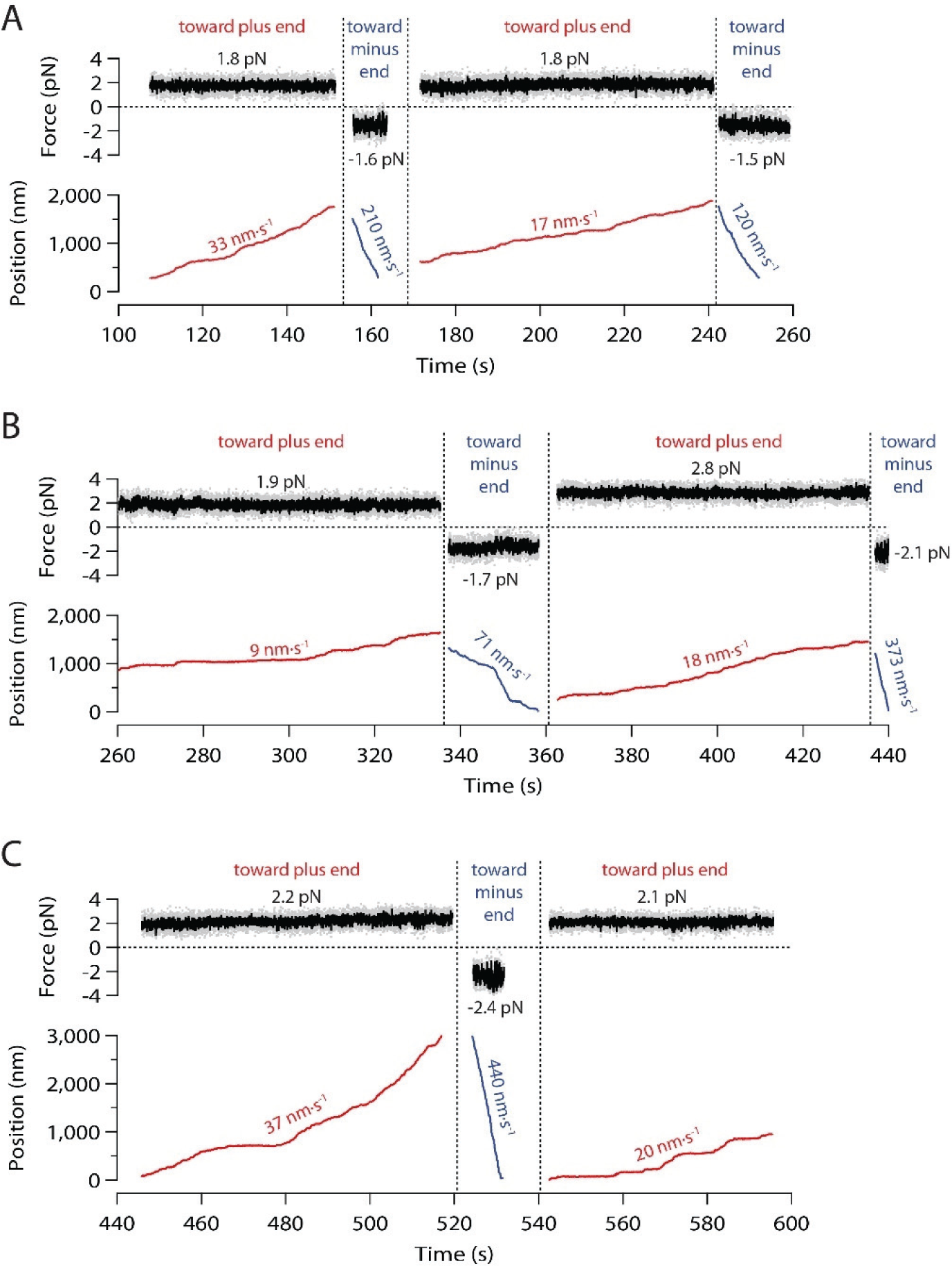
Example records showing measurement of bidirectional sliding friction. Force and position are plotted against time for a kinetochore-decorated bead attached to the side of a coverslip-anchored microtubule and pulled alternately toward the plus (red traces) and minus end (blue traces). Mean forces and speeds for each sliding event are indicated.

**Figure S5.**
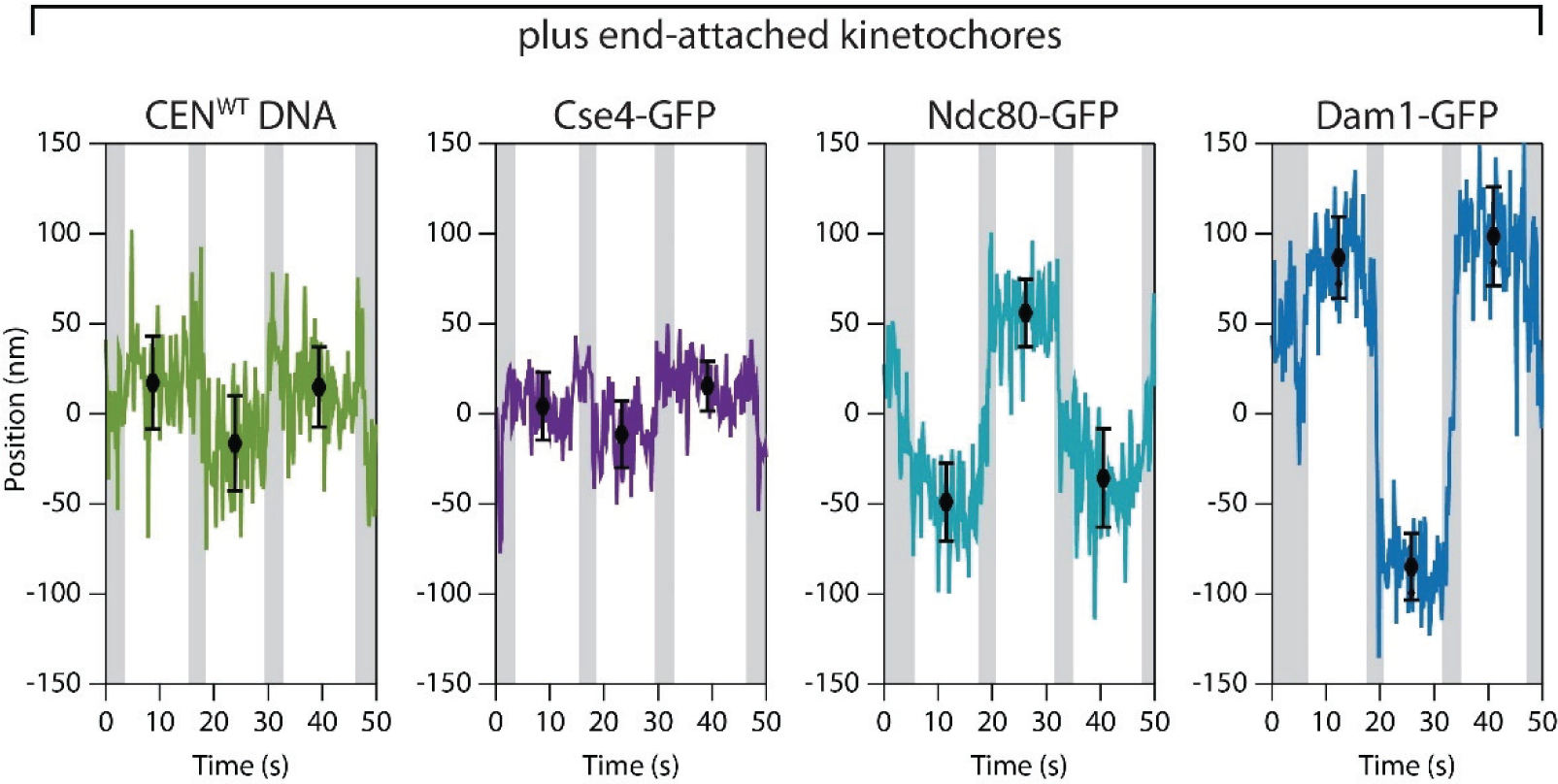
Nanoscale displacement of fluorescent-tagged components within individual plus end-attached kinetochores during periodic flow-induced reorientation. Positions for the indicated GFP-labeled components were tracked with sub-pixel accuracy while the direction of fluid flow was oscillated, causing the kinetochore and its captured microtubule to flip back and forth, reorienting by 180° with each reversal of the flow. Displacements from the tether point were estimated by averaging during the intervals when the microtubule orientation was steady. Black symbols represent mean ± SDEV from *N* ≈ 60 tracked positions during each interval. Positions recorded during reorientation of the microtubule were omitted from the averaging and are indicated here by gray shading.

